# Rapid evolution of piRNA-mediated silencing of an invading transposable element was driven by abundant *de novo* mutations

**DOI:** 10.1101/611350

**Authors:** Shuo Zhang, Erin S. Kelleher

## Abstract

The regulation of transposable element (TE) activity by small RNAs is a ubiquitous feature of germlines. However, despite the obvious benefits to the host in terms of ensuring the production of viable gametes and maintaining the integrity of the genomes they carry, it remains controversial whether TE regulation evolves adaptively. We examined the emergence and evolutionary dynamics of repressor alleles after *P*-elements invaded the *Drosophila melanogaster* genome in the mid 20^th^ century. In many animals including *Drosophila*, repressor alleles are produced by transpositional insertions into piRNA clusters, genomic regions encoding the Piwi-interacting RNAs (piRNAs) that regulate TEs. We discovered that ∼94% of recently collected isofemale lines in the *Drosophila* Genetic Reference Panel (DGRP) contain at least one *P-*element insertion in a piRNA cluster, indicating that repressor alleles are produced by *de novo* insertion at an exceptional rate. Furthermore, in our sample of ∼200 genomes, we uncovered no fewer than 80 unique *P-*element insertion alleles in at least 15 different piRNA clusters. Finally, we observe no footprint of positive selection on *P-*element insertions in piRNA clusters, suggesting that the rapid evolution of piRNA-mediated repression in *D. melanogaster* was driven primarily by mutation. Our results reveal for the first time how the unique genetic architecture of piRNA production, in which numerous piRNA clusters can encode regulatory small RNAs upon transpositional insertion, facilitates the non-adaptive rapid evolution of repression.

## INTRODUCTION

Transposable elements (TEs) are widespread genomic parasites that increase their copy number by mobilizing and self-replicating within their host genomes. TEs impose a severe mutational load on their hosts by producing deleterious insertions that disrupt functional sequences (Levis et al. 1984; McGinnis et al. 1983), causing DNA damage through encoded endonucleases (Gasior et al. 2006), and mediating ectopic recombination leading to structural rearrangements (Lim 1988). These fitness costs lead to strong purifying selection against TEs in natural populations (Charlesworth and Langley 1989; Wright et al. 2001; Boissinot et al. 2006; González et al. 2008). TE expression and proliferation are furthermore strictly regulated, particularly in germline cells where TEs are exceptionally active and resulting mutations are transmitted to offspring. In the germline of most metazoans, TEs are controlled by a conserved small RNA-mediated pathway, in which Piwi-interacting RNAs (piRNAs), in complex with Argonaute proteins, silence TEs in a sequence-specific manner (Houwing et al. 2007; Brennecke et al. 2007; Aravin et al. 2007; Girard and Hannon 2008).

On evolutionary time scales, TEs are frequently horizontally transferred between non-hybridizing species, allowing TE families to colonize new host genomes (Thomas et al. 2010; Dotto et al. 2015; Peccoud et al. 2017). Although host regulation of endogenous TEs by piRNAs is ubiquitous, how the host evolves repression to novel TEs invading the genome remains poorly understood. After invasion, repressor alleles are proposed to arise through *de novo* mutation, when an invading TE copy inserts into a piRNA producing locus referred to as a piRNA cluster (Khurana et al. 2011; Girard and Hannon 2008). The existence of numerous alternative piRNA clusters (*e.g.*, 142 loci or ∼3.5% of assembled *D. melanogaster* genome based on Brennecke et al. 2007) may facilitate the evolution of repression by increasing the mutation rate to generate repressors (Kelleher 2016; Kelleher et al. 2018; Kofler 2019). However, the technical challenge of annotating polymorphic TE insertions in repeat-rich piRNA clusters has limited the identification and study of these repressor alleles. Furthermore, for most TE families it is impossible to distinguish repressor alleles that arose via *de novo* insertion into existing piRNA clusters from the reciprocal: *de novo* piRNA clusters that arose at existing TE insertions. In particular, recent studies suggest that novel piRNA clusters may emerge frequently via epigenetic mutation, when a change in chromatin state triggers bi-directional transcription and piRNA production (de Vanssay et al. 2012; Le Thomas et al. 2014; Shpiz et al. 2014; Hermant et al. 2015).

The role of selection in the evolution of host TE repression, through piRNA-mediated silencing or otherwise, also remains controversial. In sexually reproducing organisms, the selective advantage of a repressor allele is limited by recombination, which separates the repressor from the DNA it has protected from deleterious mutation (Charlesworth and Langley 1986). Additionally, while selection for repression may be strong when the genome is invaded by a new TE family, it is unclear whether it is sustained for a sufficient number of generations to enact meaningful changes in repressor allele frequency (Lee and Langley 2012). On the other hand, forward simulation models suggest that piRNA-mediated repressor alleles are targets of positive selection, especially when transposition rates are high and TEs are highly deleterious (Lu and Clark 2010; Kofler 2019; Kelleher et al. 2018). Moreover, an early population genomic analysis of *D. melanogaster* suggests that TE insertions in piRNA clusters may segregate at higher frequency than non-cluster insertions, although this is based on modest sample size and read depth (Lu and Clark 2010).

The recent invasion of both *Drosophila melanogaster* and *Drosophila simulans* by *P*-element DNA transposons (Kidwell 1983; Anxolabéhère et al. 1988; Kofler et al. 2015) provides a unique opportunity to study not only the contributions of *de novo* mutation to the evolution of piRNA-mediated silencing by resolving the location of piRNA clusters both before and after an invasion event, but also evolutionary dynamics of repressors when selection is most strong. Unlike most TEs that have inhabited their host genome for a long evolutionary time, *P-*elements invaded the *D. melanogaster* genome around 1950 by horizontal transfer from *D. willistoni* (Daniels et al. 1990; Kidwell 1983; Anxolabéhère et al. 1988).

Subsequently, *D. simulans* acquired *P*-elements from *D. melanogaster* around 2010 (Kofler et al. 2015). In response, many natural populations of *D. melanogaster* evolved piRNA-mediated repression in less than 50 years (Jensen et al. 2008; Brennecke et al. 2008; Kidwell 1983). However, numerous strains collected prior to both invasions are retained in laboratories and stock centers, providing a historical record of ancestral piRNA clusters that were active before the *P-*element invasion.

*P-*element insertions into piRNA clusters have a well-established role in the evolved repression of *P-*elements in both *D. melanogaster* and *D. simulans*. Of particular significance are *P-*element insertions into subtelomeric piRNA clusters located in telomeric associated sequence (TAS). Seven unique wild-derived *P*-element insertions into *X*-TAS have been demonstrated to confer maternal repression of *P*-element transposition (Ronsseray et al. 1996, Marin et al. 2000, Stuart et al. 2002), three of which have been independently demonstrated to produce *P*-element derived piRNAs (Brennecke et al. 2008). Similarly, in laboratory populations of *D. simulans,* evolved repression is associated with the insertion of *P-* elements into the TAS piRNA cluster on chromosome *3R* (Kofler et al. 2018). Finally, in *D. melanogaster*, *P-*element insertions into non-TAS piRNA clusters on chromosomes 3 and 4 have also been demonstrated to confer piRNA-mediated repression (Moon et al. 2018; Khurana et al. 2011), supporting a general role for piRNA clusters in controlling *P-*elements.

Here, we take advantage of ∼200 fully sequenced *D. melanogaster* genomes, comprising the *Drosophila* Genetic Reference Panel (DGRP)(Mackay et al. 2012; Huang et al. 2014), to study the emergence and evolutionary dynamics of piRNA-mediated repressor alleles after the *P*-element invasion into *D. melanogaster* populations. To differentiate *de novo* insertions into ancestral piRNA clusters from novel piRNA clusters, we identified piRNA clusters in *D. melanogaster* from 9 *P*-element free strains of *D. melanogaster* collected before invasion. Furthermore, to empower the identification of repressor alleles, we developed a novel approach to identify TE insertions in repetitive DNA. We show that more than 90% of DGRP lines have at least one *P*-element in an ancestral piRNA cluster, with 81% containing an insertion in TAS clusters, indicating *P*-element repressors are widespread in natural populations. Moreover, we detected no fewer than 80 independent *P*-element insertions in ancestral piRNA clusters, suggesting an exceptionally high *de novo* mutation rate for the formation of piRNA-mediated repressor alleles. Finally, we found no evidence for positive selection on piRNA cluster insertions after accounting for a known insertion bias in the *X-*TAS (Karpen and Spradling 1992), suggesting that mutation alone may be sufficient to explain the rapid evolution of *P-* element repression in *D. melanogaster*.

## RESULTS

### North American strains strongly repress *P-*elements

Both recent and historic samples suggest that *P-*elements are robustly repressed in North American populations of *D. melanogaster* (Itoh et al. 2007; Kidwell 1983). To confirm that this is also true for the DGRP genomes, collected in North Carolina in 2003, we assayed *P-*element repression in dysgenic crosses between DGRP females and Harwich males. In the absence of maternally deposited piRNAs, offspring of such crosses are sterile, exhibiting atrophied ovaries (Kidwell et al. 1977; Brennecke et al. 2008; Kelleher 2016). We observed that for 97.6% DGRP maternal genotypes we sampled (41 out of 42), F1 offspring of were fertile, suggesting the presence of maternally deposited piRNAs (Fig. 1A). The single strain that did not show strong repression (DGRP531) was also very difficult to maintain in the lab, suggesting that infertility may be unrelated to *P-*element activity.

**Figure 1.**
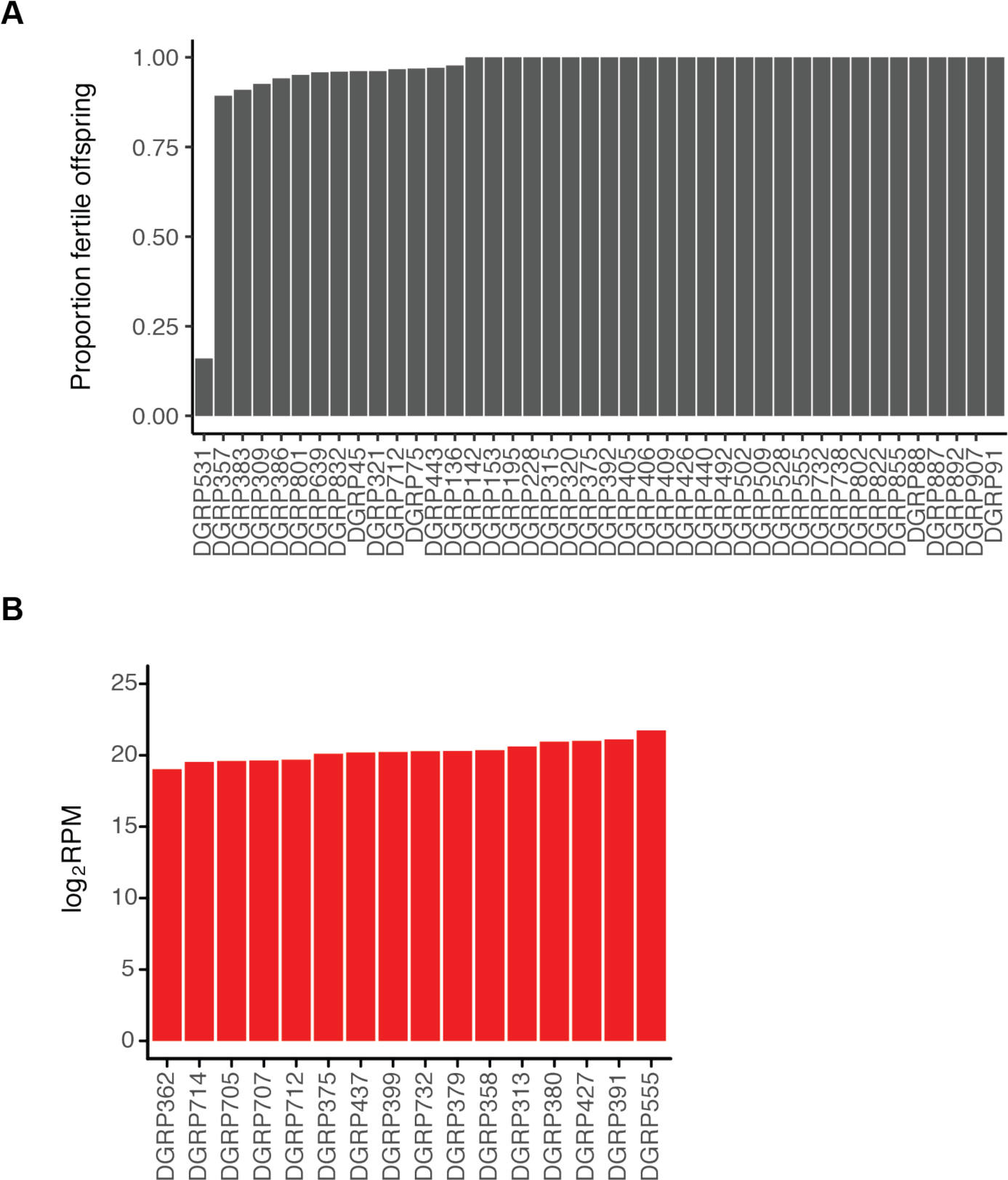
(A) Maternal repression of *P-*element activity was measured for 42 randomly selected DGRP genotypes through dysgenic crosses with males from the reference P strain Harwich. The proportion of F1 offspring from each cross is represented, with high fertility indicating maternal repression. B) *P-*element derived piRNA production for 16 DGRP genomes, based on small RNA libraries from Song *et al*. (Song et al. 2014). piRNA density is measured in Reads Per Million mapped microRNA reads (RPM) and transformed to log_2_ scale (log_2_ RPM).

We also looked directly at the production of *P-*element derived piRNAs among the DGRP using a previously published set of ovarian small RNA libraries from 16 DGRP genomes (Song et al. 2014). We discovered that all 16 produce a robust number of *P-*element derived piRNAs (Fig. 1B), consistent with the repressive phenotypes we observed (Fig. 1A). Taken together these observations suggest that maternal piRNA-mediated repression is prevalent, if not ubiquitous, among DGRP genomes.

### Identification of ancestral piRNA clusters

To uncover the genetic basis of piRNA-mediated repression, we first sought to annotate ancestral piRNA clusters in the *D. melanogaster* genome, which acted as source loci for piRNAs prior to the introduction of *P-*elements. We took advantage of 27 small RNA sequencing libraries from 9 wild-type strains (Supplemental Table S1), which were isolated from nature prior to *P*-element invasion and are therefore devoid of genomic *P*-elements. Using proTRAC (Rosenkranz and Zischler 2012), we annotated piRNA clusters based on the density of mapped piRNAs. By varying the density of reads required to annotate a piRNA cluster (pdens = 0.01, 0.05, and 0.10), we generated three sets of annotations, which contained 32, 159 and 497 piRNA clusters, and comprised 0.30%, 1.27 %, 3.68% of the assembled *D. melanogaster* genome, respectively (Fig. 2A; Supplemental Table S2, S3).

**Figure 2.**
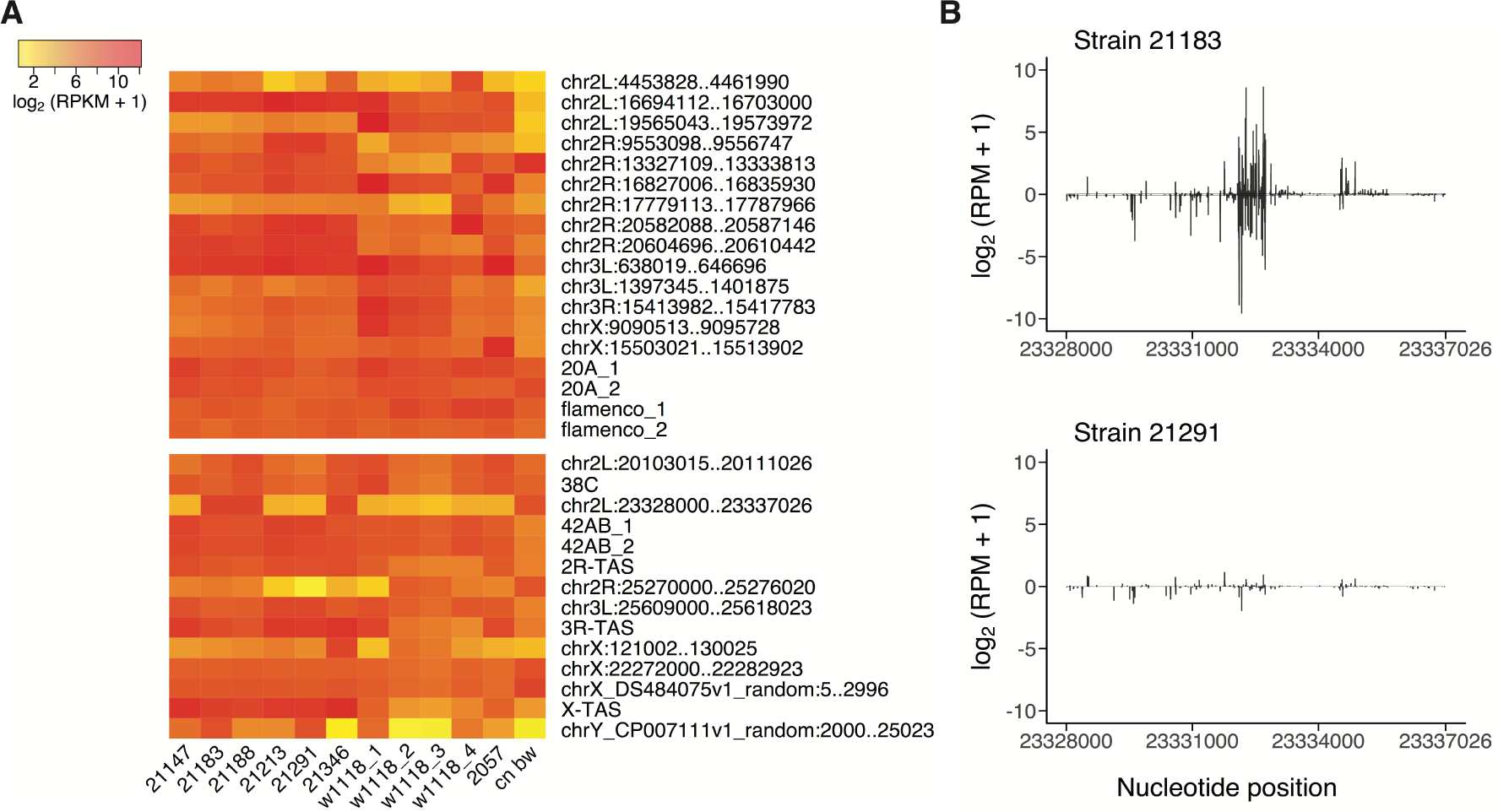
(A) Activity of 32 piRNA clusters in ancestral (*P-*element free) strains of *D. melanogaster*. Each column represents a small RNA sequencing library (biological replicates are combined) and each row represents a piRNA cluster annotated in at least one of these libraries. Coordinates of piRNA clusters are based on the *D. melanogaster* release 6 assembly (dm6: Hoskins et al. 2015). piRNA cluster expression levels are estimated by Reads Per Kilo base per Million mapped reads (RPKM) and transformed to log_2_ scale (log_2_ (RPKM + 1)). Clusters above the white line are uni-strand piRNA clusters and ones below the white line are dual-strand piRNA clusters. Details on small RNA library prep, which may be related to differences in annotated piRNA clusters between libraries from *w1118*, are provided in Supplemental Table S1. (B) An example of a polymorphism in piRNA cluster activity in an annotated cluster on chromosome 2L (23328000..23337026). Abundant piRNAs are detected from strain 21183, whereas strain 21291 produces few piRNAs. Only uniquely mapping piRNAs are considered. piRNA density is measured in Reads Per Million mapped reads (RPM) and transformed to log_2_ scale (log_2_ (RPM + 1)). Positive value represent piRNAs mapped to sense strand of the reference genome, and negative value represent piRNAs from antisense strand.

**Table 1.** *P*-element insertions in TAS

| TAS array | # of genomes with<br>insertions | # of alleles |
| --- | --- | --- |
| 2*R*-TAS | 32 | 19 (19) |
| 3*R*-TAS | 34 | 16 (16) |
| *X*-TAS | 127 | 50 (45) |
| 2*L* and 3*L*-TAS | 0 | 0 (0) |

Numbers in parentheses indicate PCR verified alleles.

We identified some genomic loci that differ in their status as a piRNA cluster between genotypes, producing abundant piRNAs in some strains while remaining quiescent in others (Fig. 2A-B; Supplemental Table S2, S3). We therefore defined ancestral piRNA clusters as genomic regions that were annotated from at least one small-RNA library. Notably, major known piRNA clusters such as *flamenco* and *42AB* (Robert et al. 2001; Malone et al. 2009; Li et al. 2009; Brennecke et al. 2007) produced abundant piRNAs in all genotypes, and were annotated as piRNA clusters regardless of stringency (Fig. 2A; Supplemental Table S2, S3). In light of clear examples of polymorphism, our annotations should not be considered a comprehensive list of the piRNA clusters segregating in ancestral populations, but rather a representative sample that includes most clusters segregating at high frequency or fixed at the time of invasion.

### Most North American Genotypes have *P* elements in ancestral piRNA clusters

To identify *P*-element insertions in extant populations, we took advantage of the published genomes from the DGRP, which includes more than 200 fully sequenced inbred lines (Mackay et al. 2012). All DGRP genomes are known to harbor *P*-elements (Zhuang et al. 2014; Rahman et al. 2015)(Supplemental Fig. S1) and the majority of them are expected to repress *P*-element activity (Kidwell et al. 1983; Anxolabéhère et al. 1988; Kidwell 1983). Although previous annotations suggest that less than 20% DGRP genomes have *P*-elements in ancestral piRNA clusters (based on 32 annotated piRNA clusters)(Zhuang et al. 2014; Rahman et al. 2015), we suspected that this was a gross underestimate, because the common requirement for unique read alignment to the reference genome prohibits the identification TE insertions in repeat-rich piRNA clusters. This is particularly problematic for identifying TE insertions in TAS clusters, which are comprised of complex satellite repeats (Yin and Lin 2007; Asif-Laidin et al. 2017; Karpen and Spradling 1992). Indeed, ∼50% of wild-derived genomes are believed to harbor a *P*-element insertion in *X*-TAS (Ajioka and Eanes 1989; Ronsseray et al. 1989; Biémont et al. 1990). In light of this unusual frequency of *P-*element insertions into *X-*TAS, and the previously well-established role of TAS insertions in the regulation of *P-* elements, we pursued alternative approaches to annotating *P*-elements from Illumina-data, with a specific goal or improving annotation in TAS regions.

First, we annotated *P*-element insertion sites throughout the genome based on high-quality alignments of split reads (mapping quality score, MAPQ ≥ 20), which are not necessarily unique, yet still support a particular genomic location with high confidence. We further removed potential false positives and false negatives based on the realignment of reads to pseudo-genomes corresponding to each proposed *P-* element insertion (see materials and methods). Including high-quality non-unique alignments increases the number of annotated *P-*elements by 71% and 66% when compared to TEMP and TIDAL, respectively, two approaches that rely on unique alignments (Zhuang et al. 2014; Rahman et al. 2015; Fig. 3A; Supplemental Table S4). While we did not validate these new insertions, six out of seven additional insertions we identified in DGRP492 were also detected by previous study using hemi-specific PCR, indicating they are true insertions (Zhang and Kelleher 2017).

**Figure 3.**
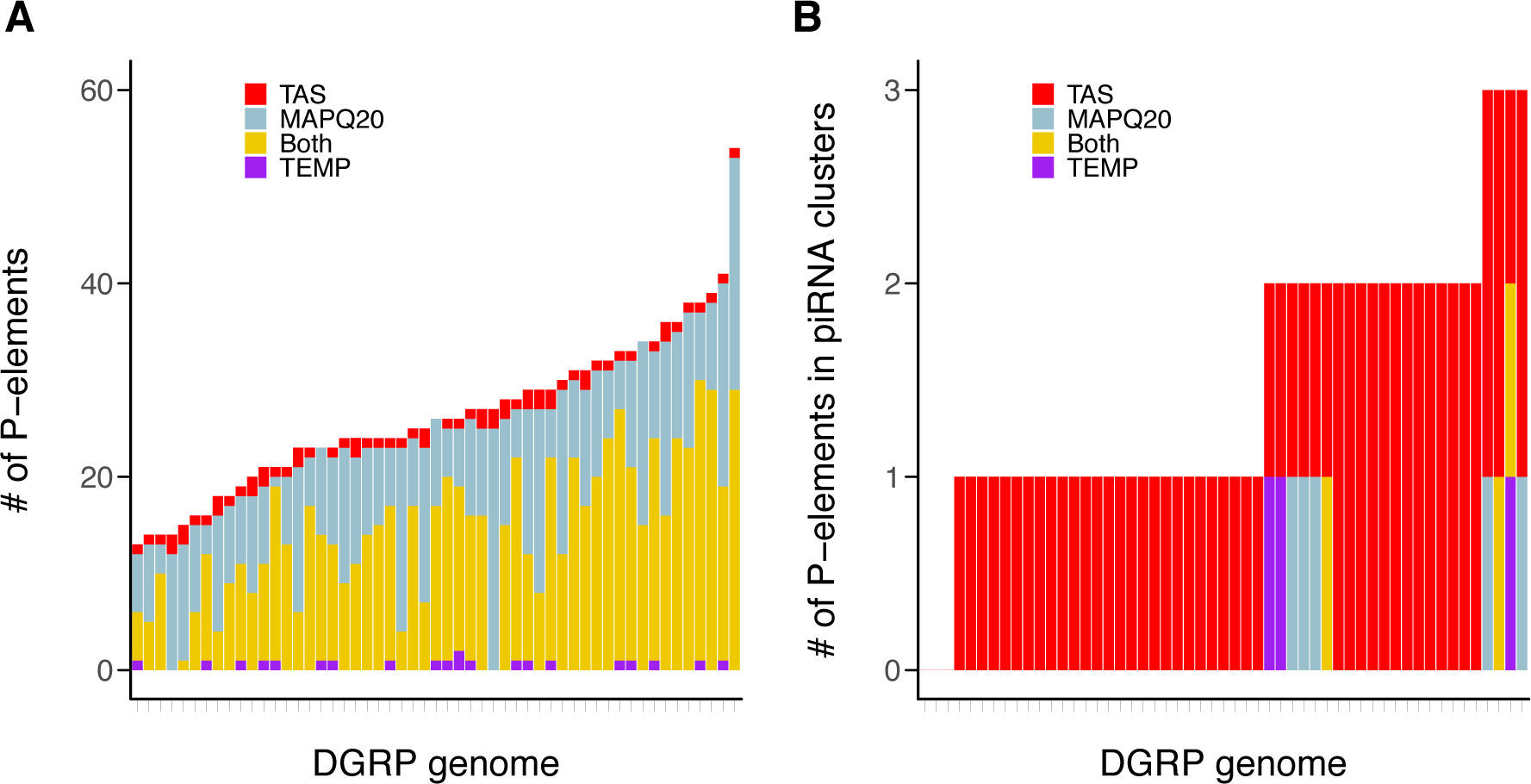
(A) Total number of *P*-elements and (B) number of *P*-elements in ancestral piRNA clusters annotated by different approaches. 53 DGRP genomes that were previously annotated by TEMP are compared. To identifying piRNA cluster insertions, the 32 cluster high confidence set was used. TEMP: insertions only found by TEMP; MA: insertions only found based on high-quality mapping; Both: insertions found by TEMP and MA; TAS: insertions only found when homologous TAS sequences were treated as a single locus.

Despite relaxing the requirement for unique alignment, we still identified only 11 DGRP genomes (5.6%) with *P*-elements in *X*-TAS. High-quality alignments likely fail to provide a unique insertion site in TAS repeats because highly similar tandem satellite sequences provide multiple equivalent alignments (Fig. 4A).

**Figure 4.**
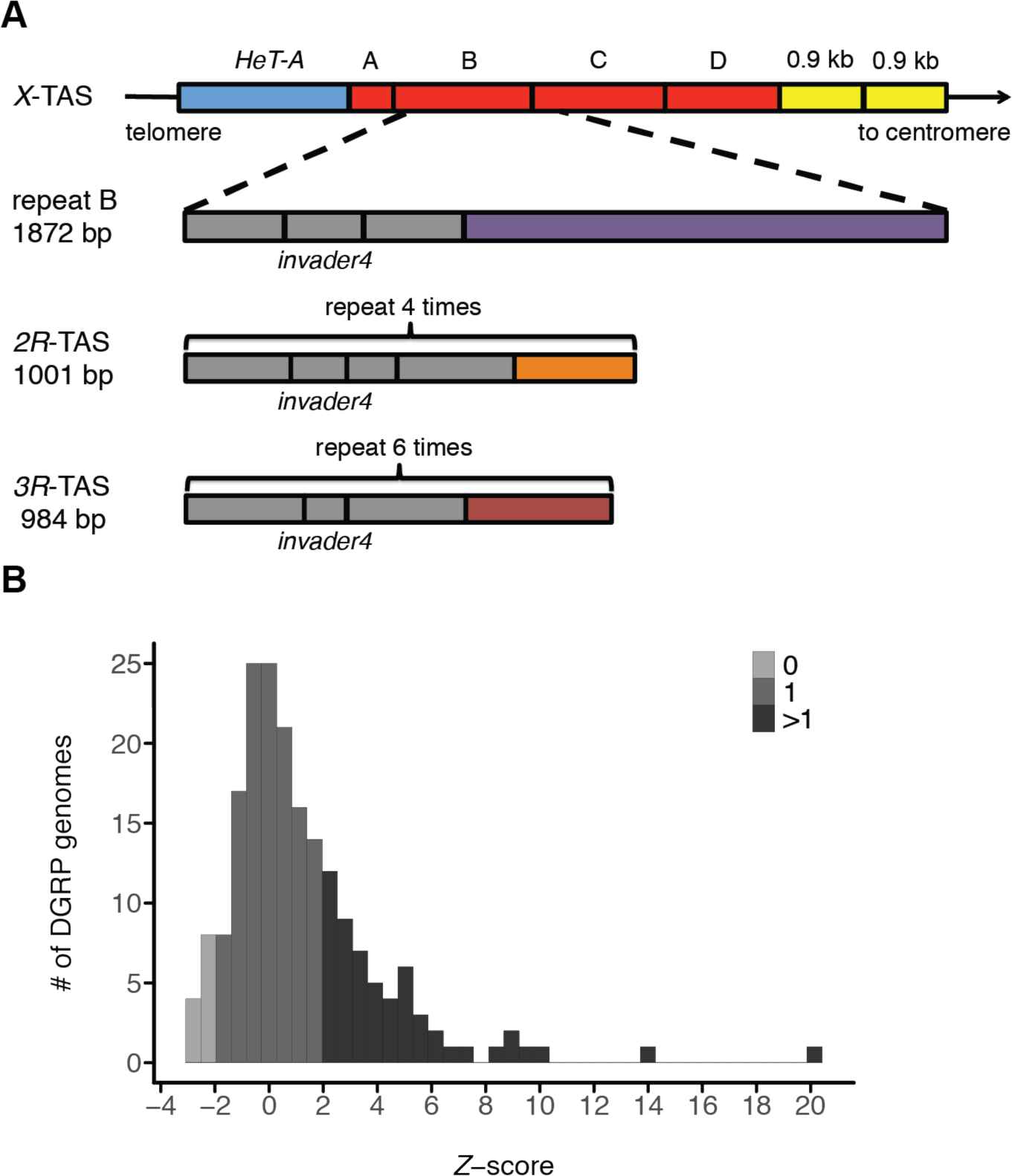
(A) The structure of TAS arrays (Asif-Laidin et al. 2017). *X-*TAS contains four tandem repeats (A, B, C and D in red) located between a *HeT-A* retrotransposon and two 0.9 kb repeats (Karpen and Spradling 1992). Repeat A is degenerated. Repeat B, C and D are ∼1.8 kb in length and highly similar to each other (>95% identity). Repeat B (enlarged below) is compared to *2R*-chromosome TAS (*2R*-TAS) and *3R*-chromosome TAS (*3R*-TAS), which are represented by 4 and 6 copies in the assembled *D. melanogaster* genome, respectively. Each repeat of *2R*, *3R* and *X*-TAS also contains several subrepeats that are composed of *invader4* retrotransposon long terminal repeats (LTRs)(gray; Bergman et al. 2006). In addition to *invader4* LTRs, the *X, 2R* and *3R-*TAS repeats share other short homologous fragments (41-131 bp) with 909 bp being unique to the *X*-TAS repeat (not shown in the figure; Asif-Laidin et al. 2017). (B) The distribution of *Z*-scores among DGRP lines. DGRP genomes with *Z* < −1.96, −1.96 < *Z* < 1.96 and *Z* > 1.96 were estimated to have 0, 1 and >1 *P*-element, respectively.

Therefore, we first sought to detect TAS insertions by identifying *P*-derived reads that aligned only to TAS repeats. We found that the majority of DGRP genomic libraries contain *P*-derived read pairs that align to *X*, *2R* or *3R*-TAS (Supplemental Table S5), while only 3 DGRP genomes contained *P*-derived reads aligning to *2L* and *3L-*TAS.

To estimate the number (0, 1, >1) of *P*-elements in *X*, *2R* and *3R*-TAS for each DGRP line, we took advantage of the distribution of the number of read pairs supporting individual insertions outside of TAS from the same genome. We then calculated a *Z*-score for the number of *P*-derived reads mapped to TAS. Using this approach we identified 12 DGRP genomes that harbor no *P-*element insertions in TAS (6%, *Z* < −1.96), 126 DGRP genomes that harbor one *P-*element insertion in TAS (65%, −1.96 < *Z* < 1.96), and 57 genomes that carry two or more insertions into TAS arrays (29%, 1.96 < *Z*) (Fig. 4B; Supplemental Table S5). Given that TAS arrays are ancestral piRNA clusters that are active in *P*-element free strains (Fig. 2A; Brennecke et al. 2007; Yin and Lin 2007), our observations reveal that the majority of DGRP genomes carry repressor alleles that arose by *de novo* insertion into existing piRNA clusters (Fig. 3B).

### Numerous repressor alleles reveal a high mutation rate to repression

We next sought to isolate individual repressor alleles that arose via *de novo* insertion into TAS arrays. First, we identified the candidate TAS array(s) containing *P-*element insertions in each DGRP genome based on proportion of *P-*derived read pairs whose best alignment supported an insertion in *X*, *2R* or *3R*-TAS (see methods; Supplemental Table S6). For the 86% of DGRP genomes in which we identified at least one candidate TAS array harboring a *P-*element insertion, we further identified the insertion site that was supported by the most read pairs (see methods). In addition, based on alternative breakpoints identified by alignments to TAS sequences, we also determined which of multiple alternate pseudo-genomes, containing *P-*element insertions into different sites, was supported by the most reads (see methods). Due to sequence homology among repeats within the same TAS array (>95% identity; Fig. 4A), we assumed all homologous insertion sites among tandem repeats corresponded to a single insertion event for these analyses.

Among 80 DGRP genomes, we found 84 *P-*element insertions into TAS where the best insertion site identified by reference genome and pseudo-genome alignments agreed, suggesting well-supported insertion sites (Supplemental Table S6). These corresponded to 40 unique insertion sites, 27 of which we were able to verify by site-specific PCR (68%). For the remaining 13 insertions, PCR revealed that seven were located at different sites, PCR failed for two sites, and PCR was not attempted for four sites. We further attempted PCR to determine the insertion sites in 22 DGRP genomes where the two computational methods did not agree, and 72 DGRP genomes where *P*-elements could not be assigned to a particular TAS or breakpoints could not be determined due to an absence of split reads. These PCRs determined an additional 43 *P-*element insertion sites in 77 DGRP genomes (Supplemental Table S6).

In total, we identified 85 independent insertions of *P-*elements into TAS sequences (*2R*, *3R* or *X*-TAS), 80 of which were verified by PCR in at least one DGRP genome (Table 1; Supplemental Table S6). Consistent with previous studies (Ajioka and Eanes 1989; Ronsseray et al. 1989; Biémont et al. 1990), we found that >50% of DGRP genomes had *P*-element insertions in *X*-TAS and ∼17% DGRP genomes had *P*-elements in *2R* and *3R*-TAS (Table 1; Supplementary Table S6). Moreover, we discovered a multitude of insertion alleles in each TAS array: 19 in *2R*-TAS, 16 in *3R*-TAS and 50 in *X*-TAS (Table 1; Fig. 5A-B; Supplemental Table S6).

**Figure 5.**
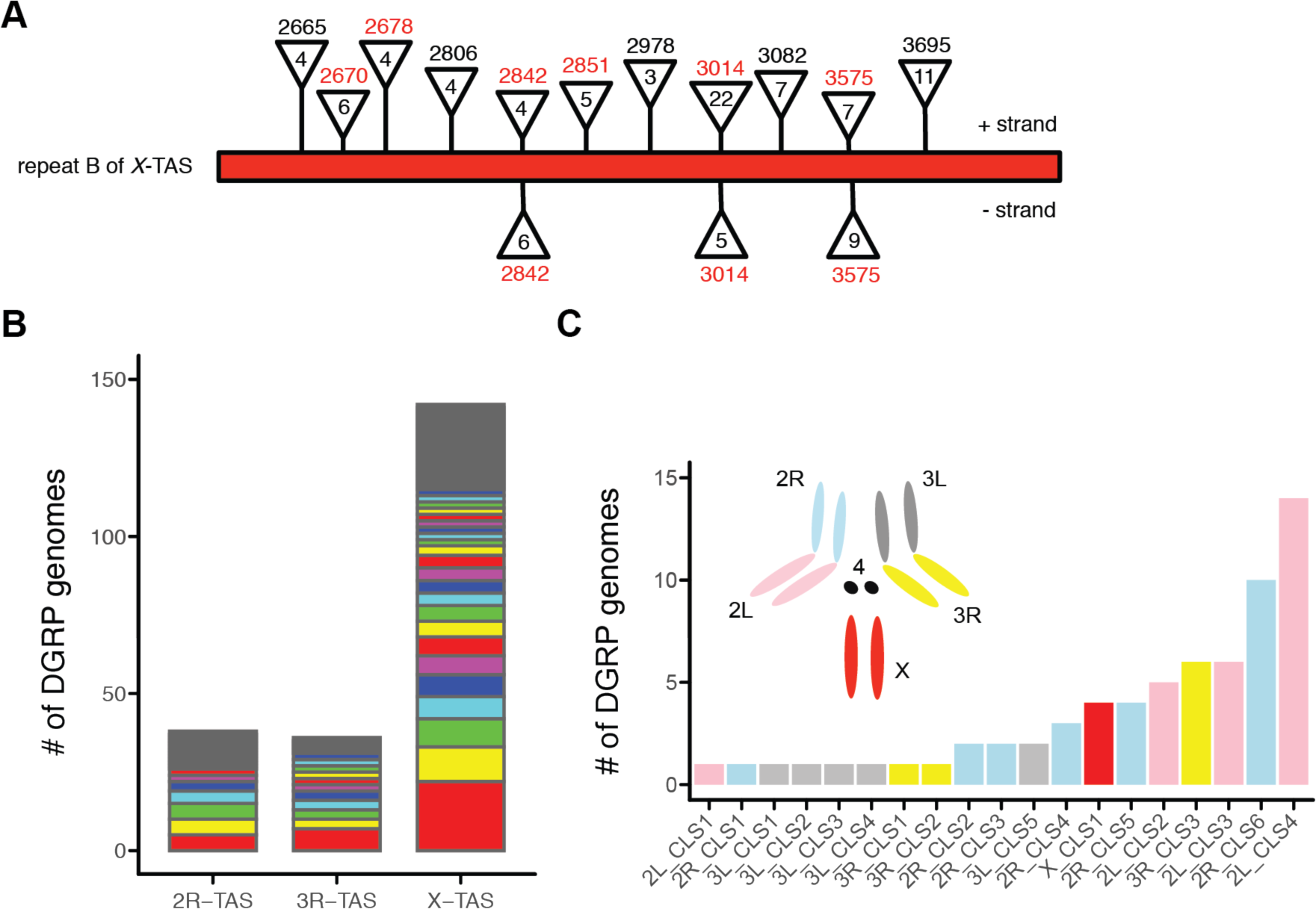
(A) Multiple *P*-element insertion sites were detected in *X*-TAS. Due to the high sequence similarity between repeats B, C and D of *X*-TAS, we arbitrarily assigned all *P*-element insertions in these repeats to repeat B. Only insertions present in more than 2 DGRP genomes are depicted. For each insertion, the breakpoint is labeled above the triangle and the number of DGRP genomes containing the insertion is indicated inside the triangle. Breakpoints positions in red correspond to insertion hotspots (Karpen and Spradling 1992). Sense *P*-elements, whose orientation is same as *X*-TAS were drawn above repeat B, whereas antisense *P*-elements were located below repeat B. (B) Multiple *P*-element insertion sites were detected in *2R*, *3R* and *X*-TAS. Each color represents a unique *P*-element insertion. (C) *P*-elements are present in non-TAS ancestral piRNA clusters across all major chromosome arms, based on our annotation set of 159 piRNA clusters.

### *P-*elements in preferred insertion sites exhibit elevate polymorphic frequencies

*P*-elements have some preferred insertion hot points, which are found in *X,* 2R and *3R-*TAS (Karpen and Spradling 1992) as well as at euchromatic sites around the genome (Spradling et al. 2011). We therefore wondered whether *P-*element insertions into these hot points were unusually common among insertion alleles from natural populations. We found that hot points were greatly enriched for *P-* element insertion alleles: 88.2% (15 out of 17) of hot points in TAS arrays had a *P*-element insertion allele, as compared to only 1.4% (55out of 3840) of non-preferred sites (Fisher’s Exact Test P-value < 10^-15^, Fig. 6A). Similarly, for non-TAS regions 57.1% (16 out of 28) of hot points have an insertion allele, when compared to 0.003% of non-preferred sites (3,642 out of 143,691,516, Fisher’s Exact Test P-value < 10^-15^, Fig. 6A). Hot points were also more likely to have two distinguishable insertion alleles, one in each strand, when compared to non-preferred sites (TAS: Fisher’s Exact Test P-value = 1.51×10^-5^; non-TAS: Fisher’s Exact Test P-value < 10^-15^; Fig. 6B).

**Figure 6.**
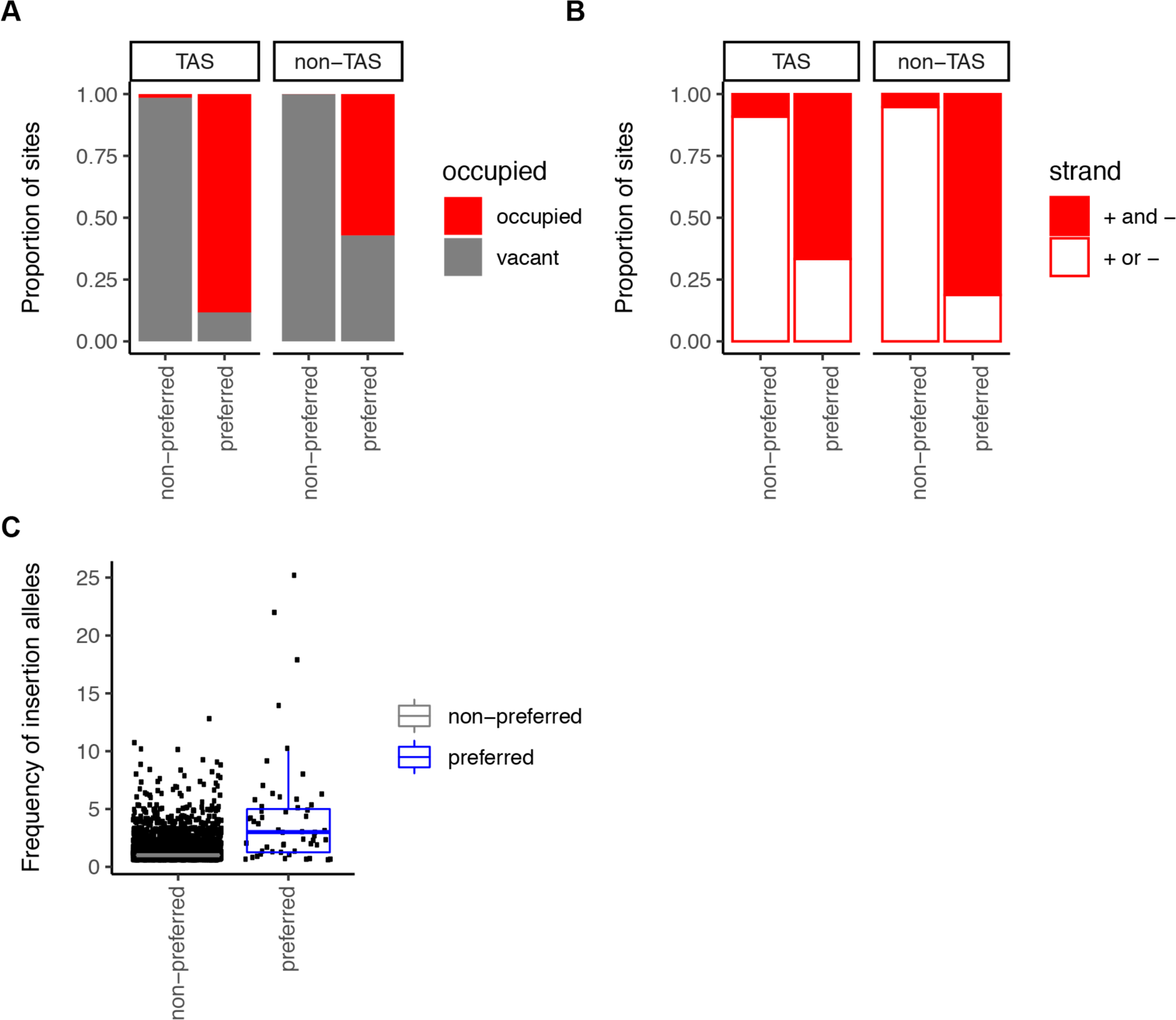
(A) The proportion of non-preferred and preferred (hot point) sites that are occupied by a *P-*element insertion in at least one DGRP genome, or vacant in all DGRP genomes. (B) The proportion of occupied sites containing *P*-element insertions in both the sense and antisense strands (red), or a single strand (white), for non-preferred and preferred sites. (C) The comparison between frequencies of *P*-elements at non-preferred and preferred sites. *P*-elements inserted at same site but in opposite orientations were considered different insertions.

Recurrent insertion into hot points potentially elevates the population frequency of insertion alleles. Indeed, even after separating insertions occurring in the same genomic position but on opposite strands, individual insertion alleles in hot points occur in a larger number of DGRP genomes than those occurring at non-preferred sites (TAS: Wilcoxon rank sum test *Z* approximation = 3.71, P-value = 2.05×10^-5^; non TAS: Wilcoxon rank sum test *Z* approximation = 8.36, P-value < 10^-15^; Fig. 6C), suggesting that recurrent insertion into these positions elevates the frequency of these insertion alleles. Taken together, our observations suggest that the exceptional abundance of *P-*element insertions in *X-*TAS arrays is at least partially explained by an insertion site preference.

### No evidence of positive selection

*P-*element insertions into ancestral piRNA clusters are proposed to benefit the host by preventing the accumulation of additional deleterious insertions, suppressing dysgenic sterility, and potentially suppressing ectopic recombination through heterochromatin formation at *P-*element loci (Charlesworth and Langley 1986; Lee and Langley 2012; Kelleher et al. 2018). Combining the TAS insertion alleles with those identified in non-TAS piRNA clusters, we detected up to 170 *P-* element insertion events into at least 15 (up to 33) different ancestral piRNA clusters, which are located on all of the major chromosome arms of the *Drosophila* genome (Fig. 5C; Supplementary Figure S2; Supplemental Table S7). Positive selection is expected to have increased the frequency of these beneficial alleles in natural populations when compared to neutral or deleterious *P-*element insertions that do not establish repression (Nielsen 2005). To test this hypothesis, we compared the frequency of insertions inside piRNA clusters to those outside of piRNA clusters among the DGRP genomes we sampled. Regardless of how stringently we defined ancestral piRNA clusters, we observed that *P-*element insertion alleles in piRNA clusters are significantly more common among DGRP genomes than those in other genomic regions (Fig. 7A-C).

**Figure 7.**
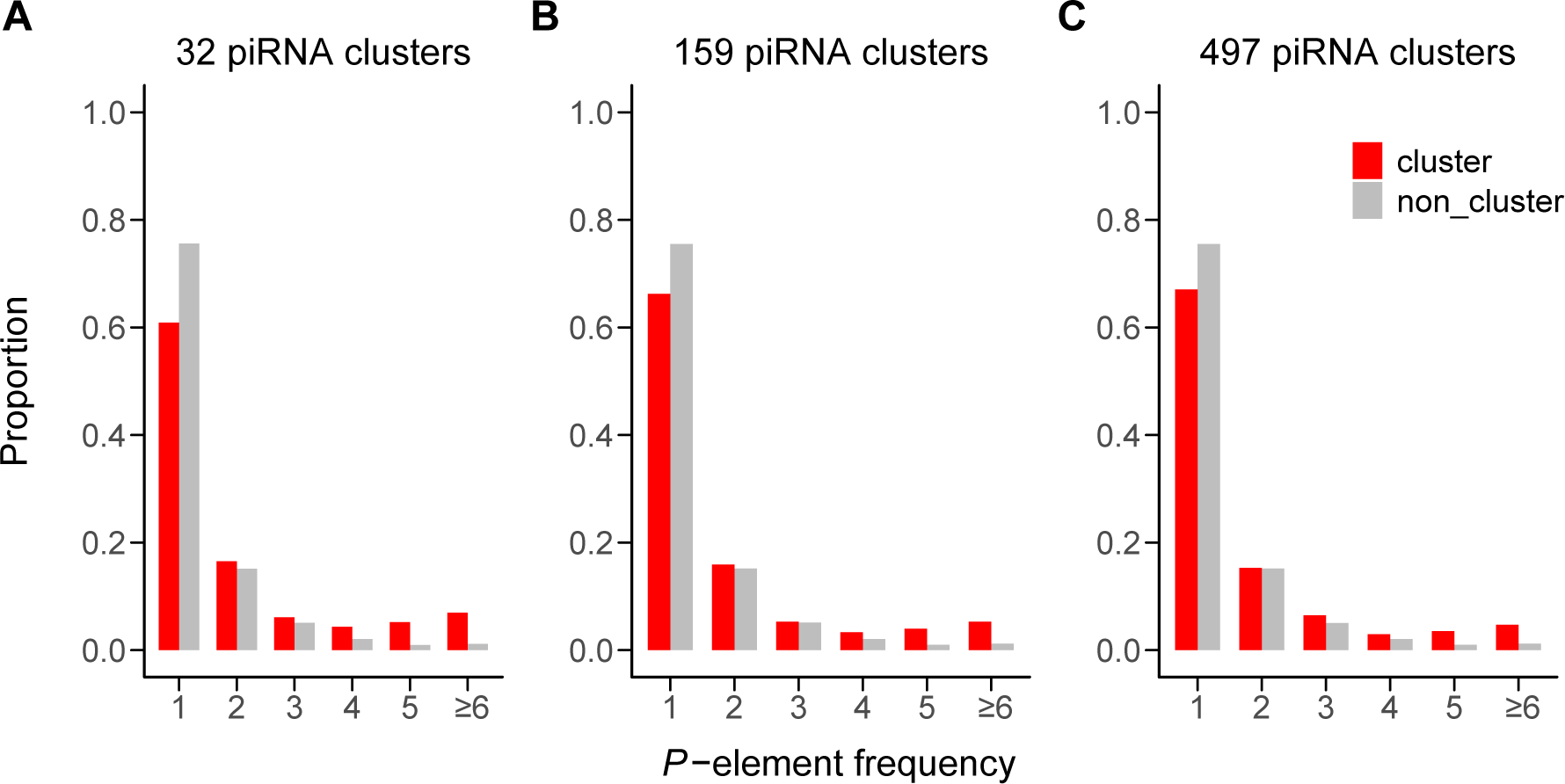
The frequency of *P*-elements in piRNA clusters (red) and the frequency of *P*-elements outside of clusters (gray) are compared for 3 sets of annotated piRNA clusters.

While the elevated frequency of cluster *P-*elements might suggest selection for repression, the observation is confounded by two factors that may also elevate the frequency of *P-*element insertions in piRNA clusters. First, recurrent insertion into hot points, which occur disproportionately in TAS piRNA clusters, elevates the frequency of those insertion alleles (Fig. 6C). Second, TE insertions rise to higher frequency in regions of low recombination, where piRNA clusters reside, most likely because of reduced purifying selection against ectopic recombination (Dolgin and Charlesworth 2008; Petrov et al. 2011; Kofler et al. 2012; Charlesworth and Langley 1989). To disentangle the potential impact of positive selection of cluster insertion from confounding effects of reduced purifying selection and recurrent insertion, we fit a multiple regression model that predicted the frequency of each *P-*element insertion as a function of its recombination rate, whether or not it occurs in a hot point, and its location inside or outside of a piRNA cluster, for all three sets of cluster annotations. Recombination rate (*F*_1,3942_ = 8.66, P-value = 0.0033) and insertion preference (*F*_1,3942_ = 404.49, P-value < 10^-15^) were both strongly associated with the polymorphic frequency of *P-*element insertion alleles. However, after accounting for these two confounding variables we were unable to detect a difference in the polymorphic frequency of insertions inside and outside of piRNA clusters (pden=0.01: *F*_1,3942_ = 0, P-value = 1; pden=0.05: *F*_1,3942_ = 0.0009, P-value = 0.98; pden=0.1: *F*_1,3942_ = 0, P-value = 1). We therefore find no evidence that positive selection has elevated from polymorphic frequencies of *P-*element insertions into ancestral piRNA clusters among DGRP genomes.

## DISCUSSION

In this study, we took advantage of the recent invasion of the *Drosophila* melanogaster genome by *P-*element DNA transposons to chronicle the evolution of piRNA-mediated repression. We reveal that the common phenotype of *P-*element repression exhibited by North American *D. melanogaster* (Ogura et al. 2007; Kidwell et al. 1983; Kidwell 1983) is underpinned by an unprecedented number of repressor alleles, which have arisen since the *P-*element invasion in the mid 20^th^ century. We uncovered no fewer than 80 unique repressor alleles, which are independent insertions of *P-*elements into piRNA clusters. Furthermore, we found no evidence that positive selection has increased the frequency of these insertions, suggesting that mutation alone is responsible for the rapid evolution of the repressive phenotype in less than ∼30 years (Kidwell 1983).

Our observations represent, to our knowledge, the first demonstration that mutation rates can be sufficiently high to drive a rapid evolutionary change. Except in cases of extreme mutation limitation, the contribution of mutation rate to the rate of evolutionary change is thought to be negligible, since the mutation rate per site is comparatively slow when compared against the action of selection. However, the exponential increase in transposition rate that occurs as TE copies accumulate, and the large numbers of functionally redundant piRNA clusters that establish repression when carrying an insertion allele, result in a mutation rate to piRNA-mediated repressor alleles that is exceptionally high (Kelleher et al. 2018). Indeed, forward simulations have previously demonstrated that piRNA-mediated repression may evolve even in the absence of any fitness cost to TEs (Kofler 2019), which mirrors the absence of a footprint of positive selection on *P-*element insertions in piRNA clusters.

*P-*element invasions may be particularly conducive to the mutation-dependent evolution of piRNA-mediated silencing, owing to the presence of multiple insertion hot points in the TAS piRNA clusters (Karpen and Spradling 1992), and their unusually high transposition rate (∼0.1 new insertions per element (Eggleston et al. 1988; Berg and Spradling 1991; Kimura and Kidwell 1994)). Indeed, 73.9% of *P-*element insertions into ancestral piRNA clusters (based on 32 annotated piRNA clusters) occurred in TAS regions, consistent with previous studies that detected *P-* elements insertions using hybridization-based approaches (Ronsseray et al. 1991; Marin et al. 2000; Stuart et al. 2002; Brennecke et al. 2008). Furthermore, we discovered that *P*-elements are most commonly observed in previously identified insertion hot points (Fig. 6A; Karpen and Spradling 1992), thereby demonstrating that this mutation bias shapes the distribution of *P-*element insertions even within the TAS clusters. TE insertions in TAS were likely not detected among DGRP genomes previously because the reliance on unique alignments excludes read pairs supporting insertions in satellite arrays (Linheiro and Bergman 2012; Zhuang et al. 2014; Rahman et al. 2015). Therefore, allowing for multiple mapping within highly homologous satellite repeats represents a powerful method for annotating TEs in these regions from short paired-end reads. Our observations echo those of a recent study of in laboratory populations of *D. simulans,* which demonstrated that *P-* element repression evolved by multiple independent insertions in piRNA clusters, particularly in the *3R*-TAS (Kofler et al. 2018).

Finally, we found that ∼94% *D. melanogaster* genomes have at least one *P*-element in an ancestral piRNA cluster, suggesting *de novo* mutation, in which *P*-elements transpose into pre-existing piRNA clusters, is the predominant mutational mechanism giving rise to piRNA-mediated silencing. Nevertheless we cannot exclude a potential role of epigenetic mutations in the evolution of piRNA-mediated *P-*element repression. Indeed, of 6 strains we examined that do not contain insertions in ancestral piRNA clusters 5 are strong repressors of *P-*element hybrid dysgenesis (Figure S3). While these strains may contain insertions into piRNA clusters that we were unable to identify (false negatives), they also may contain natural epialleles: *P-*element insertions that have been converted into heritable piRNA clusters though changes in epigenetic state (Hermant et al. 2015; Le Thomas et al. 2014; de Vanssay et al. 2012). If epigenetic mutation occurs in natural populations, it would provide an even greater increase to the already high mutation rate to piRNA-mediated silencing, further accelerating population-level transition to a repressive state.

In summary, *P*-element repression in *Drosophila melanogaster* evolved rapidly though abundant *de novo* mutations: the transposition of *P*-elements into pre-existing piRNA clusters. Strikingly, these potentially beneficial alleles exhibit no signature of positive selection, representing a heretofore-unique example of rapid evolutionary change that emerges from mutation alone. Our observations reveal how the unique genetic architecture of piRNA-mediated silencing, in which transposition into multiple functionally redundant piRNA clusters results in a repressor allele, facilitates the evolution of repression of an invading TE, thereby removing the requirement for natural selection.

## MATERIALS and METHODS

### DGRP stocks and genomes

All DGRP lines were ordered from the Bloomington Drosophila stock center.

### Assays of dysgenic sterility

Virgin DGRP females were crossed to males from the reference P strain Harwich at 29°C. 3-5 day-old F1 female offspring were assayed for ovarian development using a squash prep, as described in Srivastav and Kelleher (Srivastav and Kelleher 2017).

### piRNA cluster annotation

Ovarian small RNA sequencing libraries were downloaded from NCBI or were generated by our lab for another project (Lama and Kelleher unpublished, Supplemental Table S1). The latter libraries are available from SRA archive (SRP160954). For each library, adapters were trimmed using cutadapt (version 1.9.1) (Martin 2011). Trimmed reads with 23 – 29 nt (typical size of piRNAs in *Drosophila*) were kept for further analysis. piRNA clusters were predicted separately for each library from an ancestral *P-*element free strain using proTRAC (Rosenkranz and Zischler 2012), which identifies genomic loci corresponding to piRNA clusters based on the density of mapped piRNAs. We considered different values of the proTRAC pdens parameter (0.01, 0.05, 0.1), with lower pdens values corresponding to annotation sets that include a smaller number of higher confidence piRNA clusters. Annotated piRNA clusters less than 5 kb apart were considered a single cluster.

To identify *P-*element derived piRNAs in ovarian small RNA libraries from 16 DGRP genomes examined in Song *et al*. (2014), piRNAs were aligned to the *P-* element consensus and microRNA, respectively. *P-*element derived piRNA abundance was estimated as reads per million mapped microRNA reads (RPM).

### Detecting *P*-element insertions in DGRP genomes

DGRP whole genome sequencing reads were downloaded from the NCBI Sequence Read Archive (SRA study: SRP000694)(Mackay et al. 2012; Huang et al. 2014). 12 DGRP genomes were excluded from our analysis because 45 bp paired-end reads (DGRP357, DGRP379, DGRP427, DGRP486, DGRP786), or 75 bp single-end reads (DGRP153, DGRP237, DGRP28, DGRP325, DGRP386, DGRP41, DGRP730) were too short to allow for identification of *P-*element insertion sites. To identify read pairs that include *P*-element sequence in the remaining genomes, individual reads were separately and locally aligned to full-length *P*-element consensus (O’Hare and Rubin 1983) using bowtie2 (v2.1.0) (Langmead and Salzberg 2012) with default parameters. *P*-element sequences were then trimmed from mapped reads using a custom Perl script. Trimmed reads longer than 30 bp were kept and used for down-stream analyses. A flow chart of the annotation strategy for non-*TAS* insertions is provided in Fig. 8.

**Figure 8.**
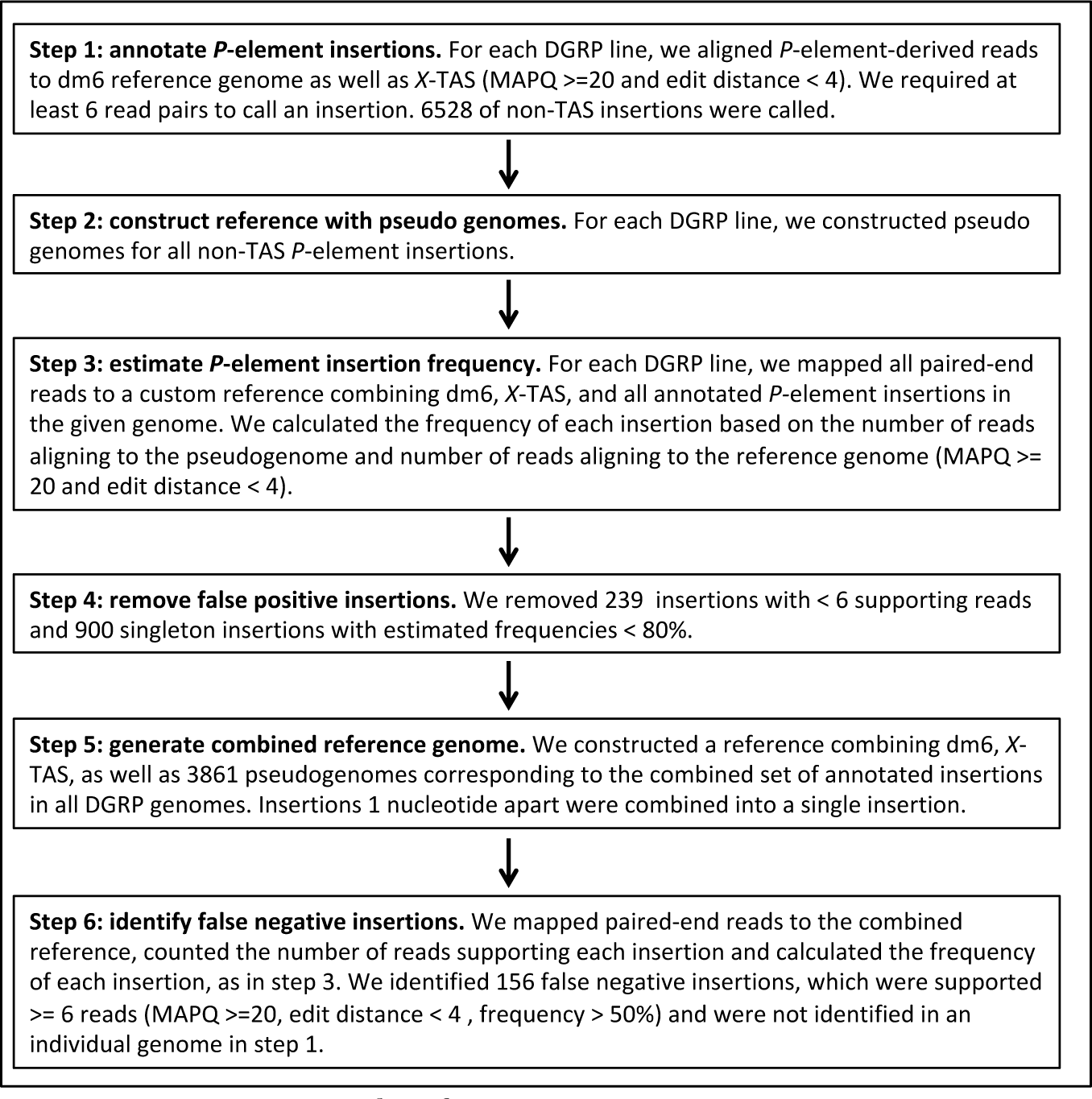
Annotation pipeline for non-TAS insertions.

For each DGRP genome, the *P*-derived trimmed reads were first aligned to the *D. melanogaster* release 6 reference genome (dm6: Hoskins et al. 2015) as well as *X*-TAS (Karpen and Spradling 1992) using bowtie2. Reported alignments with mapping quality score greater than 20 and edit distance (sum of mismatches and gaps required to convert the read sequence to the reference) less than four were kept. To isolate breakpoints corresponding to *P*-element insertion sites, we took advantage of split reads, in which one segment aligned to the *P*-element consensus and the remainder aligned to the reference genome. After breakpoints were located, all non-split *P*-derived read pairs (*i.e.* one read aligns to *P-*element, its mate to the reference genome) within 500 bp were identified. At least 6 supporting read pairs (split or non-split) were required to identify a candidate *P*-element insertion.

For each of 6528 candidate *P-*element insertions, we constructed a pseudo insertion allele containing 500 nt of genomic sequence on either side of insertions, an 8 nt target site duplication, and the full length *P-*element as consensus as in Zhang and Kelleher (2017). To identify potential false positives, all paired-end reads from each DGRP genome were re-aligned to a library of pseudo-insertions and reference alleles for all insertions sites annotated in the given genome, requiring >6 read pairs (MAPQ ≥ 20, edit distance <4). We then calculated the frequency of each insertion as the fraction of reads supporting the insertion allele. We removed 239 annotated insertions with <6 supporting reads in an individual genome, and 900 singleton insertions with estimated frequencies <80% as likely false positives.

After false positives were removed we sought to identify false negative insertions, which were not identified in a given DGRP genome due to an absence of split reads, but were annotated in another DGRP genome. To this end we constructed a combined reference genome including dm6, *X*-TAS, and pseudogenomes from the combined set of all 3861 candidate *P-*element insertions identified in any DGRP genome. Reads from all DGRP genomes were then aligned to this combined reference. We identified 156 false positive negative insertions that were supported by > 6 read-pairs from the Illumina library from a given strain (MAPQ ≥ 20, edit distance < 4), but were not annotated in our original alignments due to absence of split reads. The complete list of *P-*element insertions, including their estimated frequencies in each DGRP genome, are provided in Supplemental Table S8.

### Detecting *P*-element Insertions in TAS

We divided the dm6 reference genome into two parts: TAS and non-TAS regions. TAS regions included full-length of *X*-TAS (9872 bp, L03284) (Karpen and Spradling 1992), 2*R*-TAS (chr*2R*:25258060..25261551, 3492 bp) and *3R*-TAS (chr*3R*:32073015..32079331, 6317 bp)(Yin and Lin 2007), *2L*-TAS (chr*2L*:1..5041, 5041 bp) and *3L*-TAS (chr*3L*:1..19608, 19608 bp)(Walter et al. 1995). The other genomic regions were categories as non-TAS.

To determine if *P-*derived reads that did not map to the non-TAS regions corresponded to insertions in TAS, they were aligned to the TAS reference using bowtie2 outputting all valid alignments (-a). A read was considered mapped to TAS if the edit distance was fewer than 4. For each DGRP genome, we calculated a *Z*-score for TAS-aligned reads according to the formula: *Z* = (x – μ) / σ, where x is the number read pairs aligned to *X*, *2R* or *3R*-TAS, μ is the average number of reads supporting individual non-TAS *P*-element insertions in a given genome, and σ is the standard deviation for reads supporting non-TAS insertions. A significance level α = 0.05 (*Z* = ± 1.96) was used to estimate the number of *P*-elements in TAS in each DGRP genome (Supplemental Table S5).

To determine which TAS arrays (*X*, *2R* or *3R*-TAS) contained a *P-*element insertion in each DGRP genome, we first calculated the edit distance for all reported alignments of each read pair in that genome. We then assigned each read pair to the TAS array to which it aligned with lowest edit distance. For DGRP genomes with one *P*-element in TAS (−1.96 < *Z* < 1.96), the insertion was predicted to occur in the TAS array whose supporting reads were at least 2 times greater than the reads supporting the other two TAS arrays. For DGRP genomes with more than one *P*-element in TAS (1.96 < *Z*), we sought to determine the locations of two *P*-elements. The first insertion was predicted to occur in the TAS assay supported by the highest number of reads. Then, we subtracted the average number of reads supporting a non-TAS *P*-element insertion in the given DGRP genome from the reads supporting the first TAS insertion. The second insertion was predicted the same way as DGRP genomes with one *P*-element. The predicted *P*-element locations are provided in Supplemental Table S6.

### Localizing insertion sites of *P*-element insertions in TAS

A read pair may be equally well aligned to several homologous satellite repeats within a TAS array. Therefore, for *2R* and *3R*-TAS, we assigned *P*-elements to consensus sequences, as their repeats are indistinguishable from each other. Similarly for *X*-TAS, we were unable to determine whether a given insertion occurred in repeat B, C, or D, so we arbitrarily assigned all insertions to repeat B. We then identified the insertion breakpoint supported by the most split reads.

As an alternative approach, we also constructed pseudo genomes for each alternative TAS insertion site in a given DGRP genome, which included the *P-* element consensus sequence flanked at each end by an 8 bp target site duplication and 500 nt of adjacent TAS sequence. Paired-end reads were aligned to the constructed pseudo genomes (MAPQ ≥ 20), and the breakpoint corresponding to the pseudo genome with the most reads aligned was identified. Identified *P*-element insertion sites are provided in Supplemental Table S6.

### PCR verification of insertion sites

Genomic DNA was extracted using the QIAGEN DNeasy Blood & Tissue Kit (Cat. No. 69506) or a squish prep (Srivastav and Kelleher 2017). To determine the *P*-element insertion sites, a *P*-element specific and a TAS specific primer were used (Supplemental Table S9). As multiple bands were generally produced, owing to alternative annealing of the TAS primer to multiple repeats, the main band was purified by gel extraction using the QIAGEN MinElute Gel Extraction Kit (Cat. No. 28606), and sequenced to determine the breakpoint. PCR conditions are provided in the Supplemental Table S6.

### Recombination rates

Recombination rates at *P-*element insertions sites were identified from the genome-wide map provided by Comeron *et al*. (Comeron et al. 2012). Because these rates were based on the release 5 of *D. melanogaster* reference genome, we converted our annotated *P*-element insertions in release 6 coordinates to release 5 on the Flybase (http://flybase.org). The recombination rate of insertions that didn’t have release 5 counterparts was assumed to 0, because the major improvement of release 6 relative to release 5 is the assembly of heterochromatin regions (Dos Santos et al. 2015; Hoskins et al. 2015).

### Data analysis

Annotating piRNA clusters and identifying *P*-element insertions were powered by the high performance computing resources from the Center for Advanced Computing and Data Science (CACDS) at the University of Houston (http://www.uh.edu/cacds/resources/hpc/). All statistical analyses were performed in R (version 3.3.1)(R Core Team 2016). Graphs were made in RStudio (RStudio Team 2015) with R packages ggplot2 (version 2.2.1)(Wickham 2017b), gplots (version 3.0.1)(G. W. Warnes, B. Bolker 2016), reshape2 (version 1.4.3)(Wickham 2017a), and cowplot (version 0.7.0)(Wilke 2017).

## Acknowledgements

Shuo Zhang, Erin Kelleher, and this research were supported by a National Science Foundation Division of Environmental Biology (NSF-DEB) award to E.S.K. (NSF-DEB #1457800).

## Notes

#### Summary of Updates

We modified our TE annotation pipeline to more rigorously exclude false positive insertions. As a consequence we no longer observe that repressive P-elements, located in piRNA clusters, exhibit a signature of recent positive selection.

